# Ablation of three major phospho-sites in RyR2 preserves the global adrenergic response but creates an arrhythmogenic substrate

**DOI:** 10.1101/2024.07.08.602617

**Authors:** Jingjing Zheng, Holly C. Dooge, Héctor H. Valdivia, Francisco J. Alvarado

## Abstract

**Background:** Ryanodine receptor 2 (RyR2) is one of the first substrates undergoing phosphorylation upon catecholaminergic stimulation. Yet, the role of RyR2 phosphorylation in the adrenergic response remains debated. To date, three residues in RyR2 are known to undergo phosphorylation upon adrenergic stimulation. We generated a model of RyR2 phospho-ablation of all three canonical phospho-sites (RyR2-S2031A/S2808A/S2814A, triple phospho-mutant, TPM) to elucidate the role of phosphorylation at these residues in the adrenergic response.

**Methods:** Cardiac structure and function, cellular Ca^2+^ dynamics and electrophysiology, and RyR2 channel activity both under basal conditions and under isoproterenol (Iso) stimulation were systematically evaluated. We used echocardiography and electrocardiography in anesthetized mice, single-cell Ca^2+^ imaging and whole-cell patch clamp in isolated adult cardiomyocytes, and biochemical assays.

**Results:** Iso stimulation produced normal chronotropic and inotropic responses in TPM mice as well as an increase in the global Ca^2+^ transients in isolated cardiomyocytes. Functional studies revealed fewer Ca^2+^ sparks in permeabilized TPM myocytes, and reduced RyR2-mediated Ca^2+^ leak in intact myocytes under Iso stimulation, suggesting that the canonical sites may regulate RyR2-mediated Ca^2+^ leak. TPM mice also displayed increased propensity for arrhythmia. TPM myocytes were prone to develop early afterdepolarizations (EADs), which were abolished by chelating intracellular Ca^2+^ with EGTA, indicating that EADs require SR Ca^2+^ release. EADs were also blocked by a low concentration of tetrodotoxin, further suggesting reactivation of the sodium current (*I_Na_*) as the underlying cause.

**Conclusion:** Phosphorylation of the three canonical residues on RyR2 may not be essential for the global adrenergic responses. However, these sites play a vital role in maintaining electrical stability during catecholamine stimulation by fine-tuning RyR2-mediated Ca^2+^ leak. These findings underscore the importance of RyR2 phosphorylation and a finite diastolic Ca^2+^ leak in maintaining electrical stability during catecholamine stimulation.

## INTRODUCTION

Ryanodine receptor type 2 (RyR2) is the main Ca^2+^ release channel in cardiac muscle cells. It plays a crucial role in excitation-contraction coupling (ECC), the process that transduces electrical signals in the form of action potentials (APs) into mechanical contraction. Membrane depolarization during an AP activates sarcolemmal L-type Ca^2+^ channels (LTCC), which conduct an inward Ca^2+^ current (*I_CaL_*). *I_CaL_* then activates RyR2 channels located on the sarcoplasmic reticulum (SR), an intracellular Ca^2+^ store, triggering additional Ca^2+^ release into the cytosol (Ca^2+^-induced Ca^2+^ release or CICR). The resulting elevation of intracellular [Ca^2+^], commonly called a Ca^2+^ transient (CaT), then prompts contraction of the sarcomeres. Relaxation occurs as Ca^2+^ is removed from the cytosol by the SR Ca^2+^ pump (SERCA2) and the sarcolemmal Na^+^/Ca^2+^ ex-changer (NCX).^1^

Upon adrenergic stimulation, catecholamines bind to β1-adrenergic receptors (β1-ARs) and trigger a signaling cascade that increases heart rate (chronotropic effect), enhances contractility (inotropic effect), and expedites relaxation (lusitropic effect).^1,2^ These effects are largely mediated by downstream activation of protein kinase A (PKA) and Ca^2+^/calmodulin-dependent kinase II (CaMKII), two enzymes that regulate ion channels, as well as ECC and contractile proteins.^1,2^ In ventricular myocytes, phosphorylation of Rad mitigates its inhibition on LTCC augmenting Ca^2+^ entry via *I_CaL_*,^3^ while phosphorylation of phospholamban (PLB) reliefs its inhibition on SERCA2, boosting SR Ca^2+^ reuptake. Together, these modifications contribute to larger and faster CaTs, enhancing contractility and accelerating relaxation.^1–3^ RyR2 is the largest known ion channel and also one of the first substrates to undergo phosphorylation during adrenergic stimulation.^4^ Among the nearly 5000 residues in a RyR2 subunit, three residues are commonly recognized as part of the β1-ARs signaling and, hence, are considered “canonical”: S2031 (phosphorylated by PKA), S2808 (by PKA and CaMKII), and S2814 (by CaMKII) (human nomenclature is used throughout this report; Figure 1A).^5,6^ Despite extensive research, the specific role of the three RyR2 phosphosites in adrenergic response remains debated.^7^

**Figure 1.**
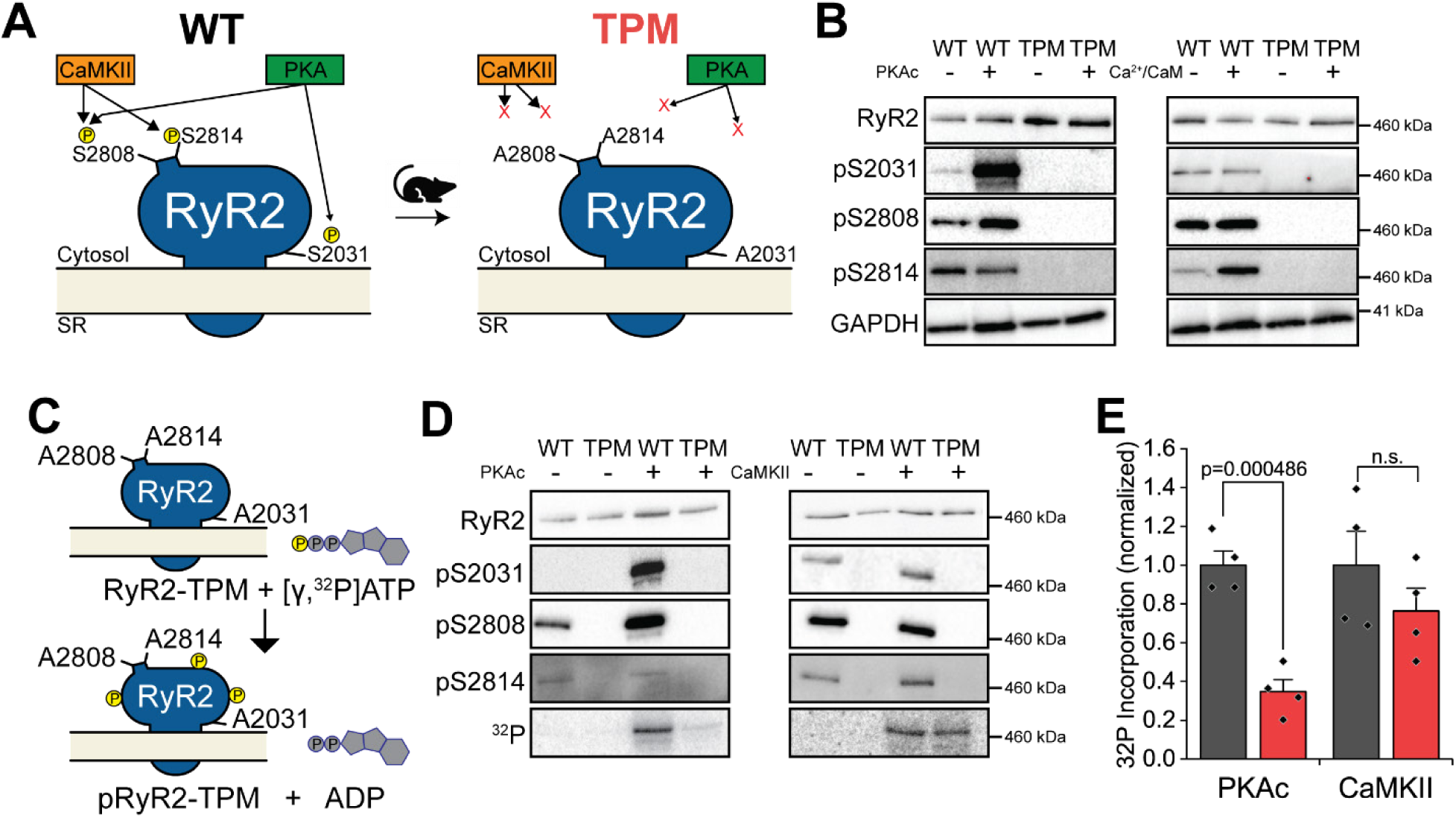
TPM channels cannot be phosphorylated at the canonical sites. **A**. Schematic diagram of RyR2 phospho-sites. Three residues in RyR2 are classically known to be phosphorylated by two kinases: S2808 and S2031 are phosphorylated by PKA, while S2808 and S2814 are phosphorylated by CaMKII. We generated a mouse model with genetic ablation of the three phosphorylation sites: RyR2-S2031A/S2808A/S2814A or triple phospho-mutant (TPM). **B**. Western blots of total RyR2 and phospho-RyR2 at the indicated sites in total heart homogenates. TPM does not show phospho-signal at the targeted residues. Addition of the catalytic subunit of PKA (PKAc) or activation of endogenous CaMKII (using Ca^2+^/CaM) leads to phosphorylation of the three sites only in the WT samples. **C.** Diagram of in vitro phosphorylation of TPM channels in the presence of [γ,^32^P]-ATP. Incorporation of ^32^P into TPM channels indicates phosphorylation of non-canonical sites. **D**. Representative images of western blots of total RyR2 and phospho-sites, and ^32^P incorporation of RyR2 immunoprecipitated from cardiac microsomes and phosphorylated in vitro with PKA and CaMKII in the presence of [γ,^32^P]-ATP. **I**. Quantification of ^32^P incorporation into WT and TPM channels. PKA incorporated significantly less ^32^P into TPM than in WT channels, while there was no significant difference in ^32^P incorporation by CaMKII (N = 4 per group, t-test)

Landmark studies initially demonstrated that mice expressing channels with ablation of S2808 (S2808A) show blunted chronotropic and inotropic responses.^8^ It was later proposed that “hyperphosphorylation” of S2808 in heart failure is an essential component of cardiac dysfunction and arrhythmia.^9^ These results are controversial because other groups, including ours, used mice with the same mutation (S2808A) but observed an intact adrenergic response and progression to heart failure.^10,11^ Phosphorylation of S2814 by CaMKII is widely accepted to increase channel activity in multiple settings of heart disease.^7,12^ Mice with genetic ablation of this site (S2814A) show decreased RyR2-mediated SR Ca^2+^ leak, reduced propensity to arrhythmia and blunted progression to heart failure.^13^ Intriguingly, S2808 and S2814 are located in a flexible loop that may contain other phospho-sites.^14^ S2031 has gained attention recently as it was proposed to be necessary for a full adrenergic response because cardiomyocytes from mice harboring S2031A channels displayed fewer Ca^2+^ sparks and blunted ECC gain in response to adrenergic activation.^15^ Increased phosphorylation of S2031 has also been linked to arrhythmogenesis in heart disease.^16^ Finally, mice with double ablation of S2808 and S2814 (S2808A/S2814A) showed normal adrenergic response but increased propensity to arrhythmia.^17,18^ While some aspects of this overall framework are disputed, the consensus in the field is that increased phosphorylation of RyR2 increases channel activity. Since SR Ca^2+^ leak mediated by hyperactive RyR2 channels promotes cardiac arrhythmia and may contribute to cardiac dysfunction,^19^ an ensuing conclusion is that phosphorylation of one, two, or all three of these canonical sites is responsible for the RyR2 hyperactivity and deleterious Ca^2+^ leak. However, a critical confounding factor in all the studies that have addressed this question is that ablation of one canonical phospho-site site may lead to compensatory changes in the phosphorylation state of the remaining sites. For example, S2031A mice exhibited an increased basal phosphorylation of S2808 and S2814,^15^ making it difficult to ascertain whether the blunted adrenergic response was due to the lack of S2031 or the basal “hyperphosphorylated” state of S2808 and S2814.

In this study, we generated a mouse model with constitutive removal of all three canonical phosphorylation sites in RyR2: RyR2-S2031A/S2808A/S2814A (triple phospho-mutant or TPM). This model allowed us to carefully dissect the role of these residues in the regulation of RyR2 and Ca^2+^ homeostasis, while preventing potential compensatory changes in their phosphorylation levels. Remarkably, we did not observe differences in the whole heart or cellular CaT responses to adrenergic stimulation, suggesting these sites have a limited role in regulating chronotropy or inotropy. Instead, our data indicate that preventing phosphorylation of the three canonical sites results in a loss-of-function that constrains RyR2 activity, leading to reduced SR Ca^2+^ leak. Paradoxically, TPM mice showed increased propensity to ventricular arrhythmia, even though the incidence of diastolic arrhythmogenic Ca^2+^ waves and delayed after-depolarizations (DADs) was reduced in TPM myocytes. We observed faster phase-1 action potential (AP) repolarization and increased propensity for early after-depolarizations (EADs) in TPM myocytes. Therefore, our results suggest that phosphorylation of the channel may be necessary to tune subcellular SR Ca^2+^ leak. SR Ca^2+^ leak has a negative connotation because of its well-defined role in arrhythmogenesis. We surmise that a finite leak is required to maintain the electrical stability and Ca^2+^ homeostasis of cardiomyocytes during catecholamine stimulation.

## METHODS

All animal experiments received approval from the University of Wisconsin-Madison School of Medicine and Public Health Institutional Animal Care and Use Committee (protocol M5944). TPM mice were created using CRISPR/Cas9 to introduce the S2031A mutation into exon 40 of *Ryr2* in mice expressing RyR2-S2808A/S2814A channels (Figure S1).^17^ The presence of all mutations was validated by Sanger sequencing and Western blot analysis using phospho-specific antibodies. Cardiac structure and function were evaluated using echocardiography and electrocardiography (ECG) under anesthesia. Adrenergic stimulation was induced by intraperitoneal (i.p.) injection of isoproterenol (Iso, 2mg/kg) in vivo, or 300 nM Iso treatment for isolated cardiomyocytes. Catecholaminergic arrhythmia challenges in mice were performed through i.p. injection of a cocktail containing 2 mg/kg epinephrine and 120 mg/kg caffeine. Intracellular Ca^2+^ dynamics in isolated myocytes were assessed using confocal microscopy, the membrane permeable dye fluo-4 AM and field stimulation. Simultaneous membrane potential and Ca^2+^ transient (Vm/CaT) recordings, along with *I_CaL_* measurements, were obtained via whole-cell patch clamp by adding the membrane impermeable fluo-4 pentapotassium salt to the internal pipette solution. Biochemical assays, including [^3^H]ryanodine binding assays and in vitro phosphorylation reactions with [γ,^32^P]-ATP, were conducted using whole heart homogenates or cardiac SR microsomes. 129S1/SvImJ mice (Jax stock No. 002448) were used as wild type controls. Age-matched mice of both sexes, ranging from 12-20 weeks old, were used for all experiments. Statistical analysis was performed using appropriate methods (as noted in the figure and table) to compare results between TPM and WT mice. p < 0.05 was considered statistically significant. Further details are available in the Supplemental Material. Additional information is available from the corresponding author upon reasonable request.

## RESULTS

### TPM channels can undergo PKA- and CaMKII-mediated phosphorylation

We generated mice with ablation of the three major phosphorylation sites in RyR2 (S2031, S2808 and S2814, as per the human nomenclature used throughout this report) by substituting the amino acid serine (S) with the amino acid alanine (A) at codons 2030, 2807 and 2813 of the mouse *Ryr2* (Figure 1, Table S1). The resulting RyR2-S2031A/S2808A/S2814A (triple phosphomutant, TPM) mice were validated both genetically and at the protein level. Sanger sequencing confirmed the expected missense mutations at the correct positions, as well as multiple silent mutations introduced to differentiate the mutant and wild type alleles during genotyping (Figure S1B). Western blot (WB) analysis using phospho-specific antibodies demonstrated that the targeted sites do not show basal phosphorylation and are not subject to phosphorylation by specific kinases. As expected in WT samples, PKA phosphorylated residues S2031 and S2808, while CaMKII phosphorylated S2808 and S2814 (Figure 1B).^6^

Previous studies have reported that RyR2-S2808A channels cannot be phosphorylated by PKA;^9^ however, S2030 was later demonstrated to be a major substrate of this enzyme.^20,21^ Similarly, studies have shown that S2814A channels cannot undergo CaMKII-mediated phosphorylation;^12^ yet, other evidence suggests that CaMKII may target multiple residues,^22,23^ including S2808. We took advantage of the TPM channels to address these contrasting results by conducting in vitro back phosphorylation experiments in the presence of radioactively labeled ATP ([γ,^32^P]ATP) on RyR2 channels immunoprecipitated from SR microsomes. ^32^P incorporation into RyR2 can then be correlated with phosphorylation of available residues. Furthermore, ^32^P incorporation into TPM channels will indicate phosphorylation of non-canonical sites (Figure 1C).

Western blots (WBs) using phospho-specific antibodies revealed an increase in the phosphorylation of S2031 and S2808 in WT channels treated with PKA, and phosphorylation of S2808 and S2814 in WT channels treated with CaMKII (Figure 1D). As expected, TPM channels showed no signal with these phospho-specific antibodies. Notably, ^32^P incorporation was observed in both WT and TPM RyR2 channels (Figure 1D,E). The PKA-induced ^32^P signal in the TPM samples was approximately 35±6% of that observed in the WT samples (p = 0.000486, n = 4 per genotype, t-test), suggesting that S2030 and S2808 are indeed major, but not exclusive, PKA phosphorylation sites in RyR2. The CaMKII-induced ^32^P incorporation was not different with the TPM channels showing 76±12% of that of WT channels (p = 0.303, n = 4 per genotype, t-test). This suggests that S2808 and S2814 are not exclusive CaMKII phosphorylation sites. These data provide experimental evidence for the existence of additional PKA and CaMKII sites in RyR2, as previously suggested,^22,23^ although their exact molecular identities and physiological relevance remain to be established.

### Global adrenergic response is preserved in the whole heart and cellular levels

Some studies have suggested that phosphorylation of RyR2 is critical to achieve a complete cardiac adrenergic response,^15,24^ but the role of specific residues has been disputed because of contradictory data.^7^ We used a multi-level approach to assess the impact of the TPM modification on the whole heart and cellular responses to catecholamine stimulation. TPM mice developed normally and did not display an overt cardiac phenotype compared to WT controls (Figure S2A,B). Further analysis revealed no significant differences in body weight or heart weight (Figure S2C,D). Cardiac structure and function, assessed by echocardiography, were comparable between the groups, including heart rate, ejection fraction, fractional shortening, posterior wall thickness, left ventricle diameter, stroke volume and left ventricle (LV) mass (Figure S2, Table S3). The expression levels of major ECC proteins (RyR2, LTCC, NCX, and SERCA2) were not different in TPM and WT heart homogenates (Figure S3).

We evaluated the whole heart in vivo response to isoproterenol (Iso) treatment (2 mg/kg i.p.), a non-specific β-AR agonist. We employed surface ECG recordings to monitor the chronotropic response to Iso over a 12-minute period. Both TPM and WT mice showed comparable chronotropic responses, reaching peak heart rates at approximately 1-minute post-injection followed stabilization after 2 minutes (Figure 2A). Using echocardiography, we evaluated chronotropy and inotropy in basal conditions and 2 minutes after Iso injection. Both WT and TPM mice showed a similar increase in heart rate (Figure 2B), ejection fraction (Figure 2C), and fractional shortening (Figure 2D). These data suggest that TPM mice have a normal whole heart chronotropic and inotropic responses to adrenergic stimulation.

**Figure 2.**
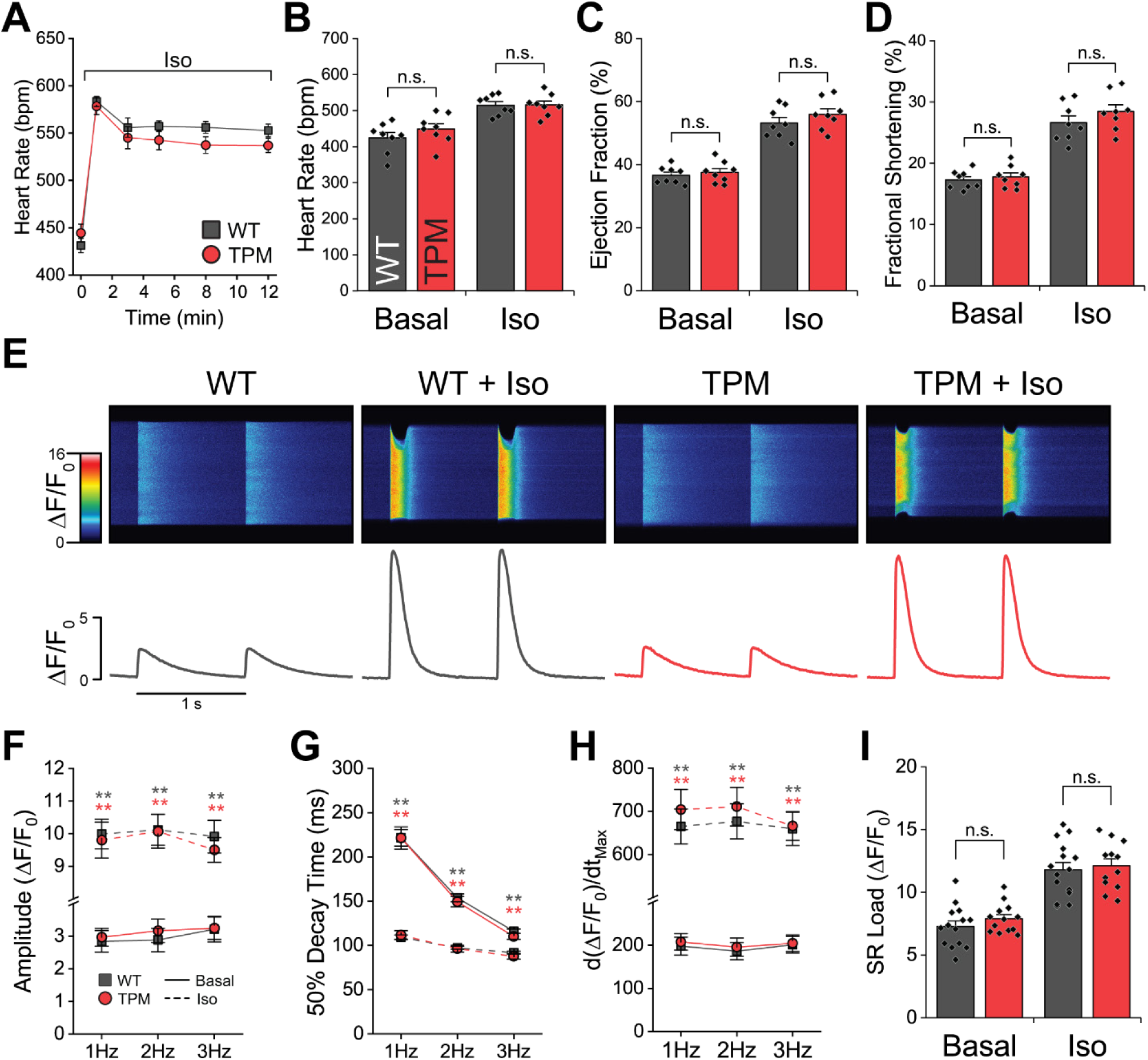
Chronotropic and inotropic responses are preserved in the TPM at the whole animal and cardiomyocyte levels. **A**. Time course of the chronotropic response to Iso stimulation (2 mg/kg i.p.) recorded by ECG in anesthetized mice. Both genotypes show a similar response without significant differences (n = 10 per genotype. Two-way repeated measures ANOVA, Holm-Sidak method). **B-D**. Chronotropic (B) and inotropic (C,D) response recorded by echocardiography in anesthetized mice 2 min after Iso injection (2 mg/kg i.p.). Both WT and TPM show a significant and comparable response to Iso. No differences were observed between genotypes (N = 8 per genotype, two-way repeated measures ANOVA, Holm-Sidak method). **E.** Representative confocal CaT images and quantified fluorescence profiles in ventricular cardiomyocytes under 1 Hz pacing in basal conditions and after 2 min of Iso perfusion (300 nM). **F-H**. Quantification of CaT properties. Both genotypes showed a comparable response to Iso stimulation in the amplitude (F), 50% decay time (G) and maximum rate of fluorescence increase (H) of the CaT, without significant differences between them at any frequency. (N = 4 hearts per genotype; n = 24 WT, 25 TPM cells [Basal]; 15 WT, 15 TPM cells [Iso]. **p < 0.01, compared to the same genotype in basal conditions. ANOVA on Ranks, Dunn’s method). **I**. SR load measured as the caffeine-induced CaT. Iso significantly increased SR load in both genotypes. No significant differences were detected between WT and TPM. (N = 4 hearts per genotype; n = 14 WT, 14 TPM myocytes [Basal]; 15 WT, 12 TPM myocytes [Iso]. Two-way ANOVA, Holm-Sidak method).

Next, we used confocal microscopy to measure Ca^2+^ transients (CaT) in isolated cardiomyocytes and evaluate the whole cell response to adrenergic stimulation. Cardiomyocytes were first paced at 1 Hz for 30 seconds and then exposed to 300 nM Iso for 2 minutes to achieve stable Ca^2+^ handling. Remarkably, the time course of the response was nearly indistinguishable between genotypes (Figure S4). A detailed assessment of rate-dependent Ca^2+^ handling was also carried out at 1-3 Hz pacing frequencies. Representative recordings at 1 Hz are shown in Figure 2E. Adrenergic stimulation caused a robust increase in CaT amplitude (Figure 2F), an acceleration of the CaT decay (Figure 2G), and an increase in the maximum rate of fluorescence upstroke (d(ΔF/F_0_)/dt_Max_, Figure 2H); yet, there were no significant differences detected between genotypes. A pulse of 10 mM caffeine applied at the end of the protocol further showed that WT and TPM myocytes have the same SR Ca^2+^ load in basal conditions and under Iso stimulation (Figure 2I). These findings suggest that the ablation of the three major phosphorylation sites in RyR2 does not affect the global Ca^2+^ transients.

### TPM channels show lower activity than WT channels

The unexpected lack of impact of the TPM modification on the whole heart chronotropic and inotropic responses or global CaT dynamics led us to further investigate the intriguing roles of the three canonical phosphorylation sites in the regulation of RyR2 channel function and sub-cellular Ca^2+^ handling. To evaluate RyR2 function at the molecular level, we performed [^3^H]ryanodine binding assays in whole heart homogenate over a range of free [Ca^2+^] (10 nM – 10 µM). Ryanodine binds to the open state of RyR2. Hence, [^3^H]ryanodine binding experiments conducted in conditions where Ca^2+^ is the only relevant activator of RyR2 can be used to probe the basal activity of the channel. The binding curves of the WT and TPM groups were nearly identical when normalized to total RyR2 protein expression (Figure 3A) or to the binding at 10 μM [Ca^2+^] (Figure 3B). As a result, TPM and WT RyR2 channels had similar maximum binding (Figure 3C) and EC50 (Figure 3D). We only noted a minor but significant decrease in [^3^H]ryanodine binding at pCa 6.5 (0.3 μM) in TPM channels compared to the WT controls (8.0±1.0% vs. 11.0±1.0%; p = 0.0289, t-test; Figure 3B).

**Figure 3.**
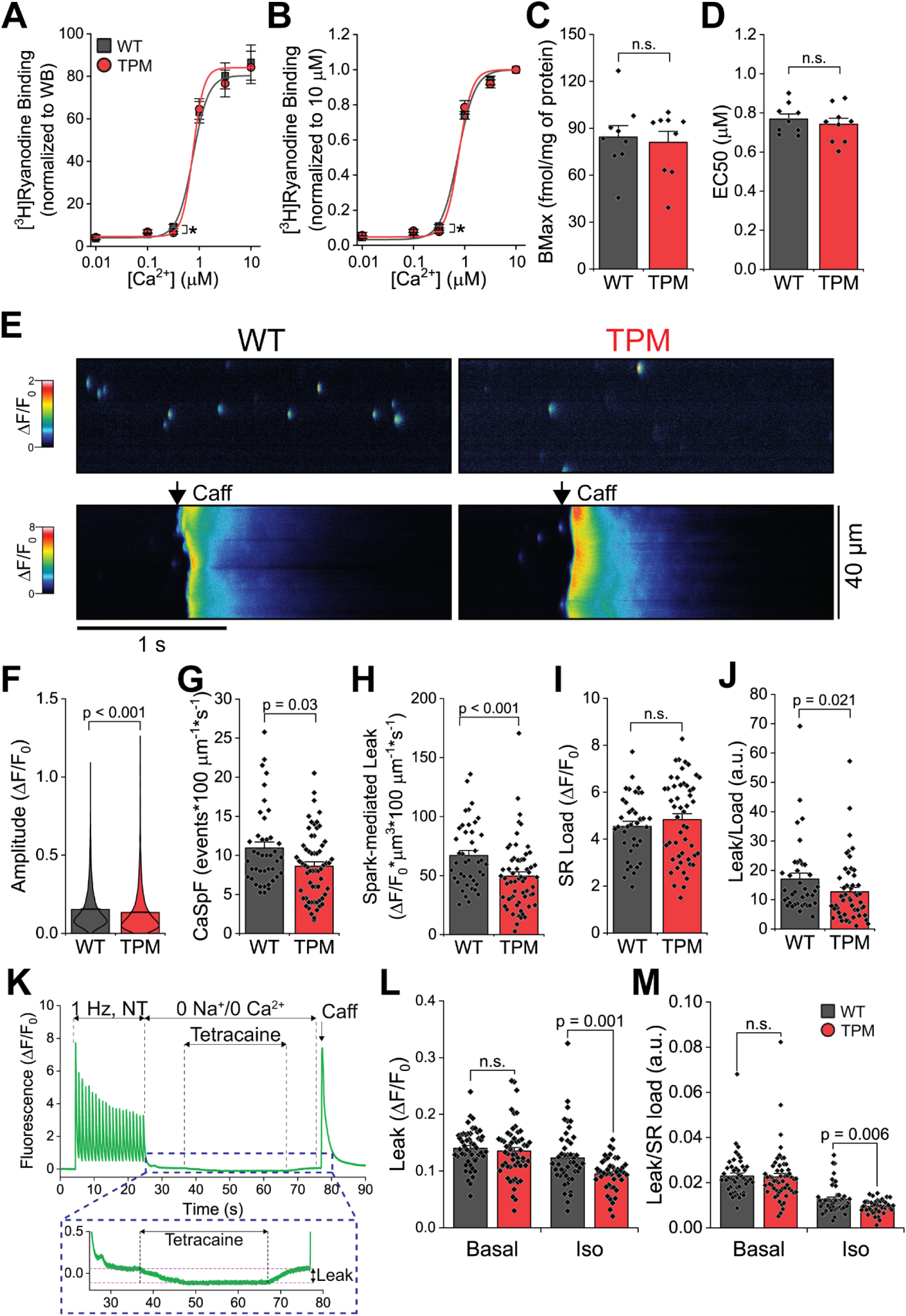
TPM myocytes show reduced RyR2-mediated SR Ca^2+^ leak. **A-B.** [^3^H]Ryanodine binding as a function of free [Ca^2+^] in heart homogenates from WT and TPM samples. [^3^H]Ryanodine binding curves were normalized to RyR2 expression from western blot data (A) and normalized to 10 µM [Ca^2+^] (B). WT and TPM RyR2 channels have similar Ca^2+^ activation profile. **C.** Average maximum binding (BMax) calculated from sigmoidal fitting of the curves in panel A. No significant difference between genotypes. **D.** [Ca^2+^] that produces 50% of maximum [^3^H]ryanodine binding (EC_50_), determined from sigmoidal fitting of the curves in panel B. No significant difference between genotypes (N = 9 per genotype. *p < 0.05. t-test or Mann-Whitney rank sum test [A-D]). **E**. Representative line scans confocal images of Ca^2+^ sparks from permeabilized cardiomyocytes. **F.** Quantification of Ca^2+^ spark amplitude. TPM sparks have significantly lower amplitude than the WT. **G.** Ca^2+^ sparks frequency (CaSpF) determined as the number of sparks in one second within a 100 μm line. CaSpF is significantly lower in TPM cardiomyocytes compared to WT. **H.** Quantification of the spark-mediated SR Ca^2+^ leak, calculated as the sum of mass from all sparks. Spark mass was calculated as previously described.^47^ Ca^2+^ leak is significantly lower in TPM cardiomyocytes (N = 4 WT, 6 TPM mice; n = 1831 sparks from 42 WT, 2101 sparks from 61 TPM myocytes. Mann-Whitney rank sum test [F-H]). **I.** SR load in permeabilized cardiomyocytes measured during acute application of 10 mM caffeine (Caff). No significant difference between genotypes. **J.** Spark-mediated leak normalized to SR load (N = 4 WT, 6 TPM mice; n = 38 WT, 51 TPM myocytes. Mann-Whitney rank sum test). TPM cardiomyocytes show reduced normalized leak. **K.** Representative fluorescence profile obtained during assessment of SR Ca^2+^ leak using the protocol by Shannon et al.^48^. Following a train of stimulation in normal Tyrode solution (NT), cardiomyocytes were perfused with 0Na^+^/0Ca^2+^ solution. Then, tetracaine (1 mM) was added to block RyR2. The bottom panel shows the enlarged area in the dashed blue box. The fluorescence drop recorded during tetracaine application reflects RyR2-mediated Ca^2+^ leak. **L-M.** RyR2-mediated SR Ca^2+^ leak (L and leak normalized to SR Ca^2+^ load (M) under basal conditions and during 300 nM Iso stimulation (N = 6 mice per genotype. n = 49-51 WT, 55 TPM myocytes [Basal]; 46 WT, 43-44 TPM myocytes [Iso]. Two Way ANOVA on Ranks (Holm-Sidak method).

At the subcellular level, we measured Ca^2+^ sparks in permeabilized myocytes with free cytosolic [Ca^2+^] clamped at 50 nM (Figure 3E). TPM cells showed Ca^2+^ sparks of lower amplitude (Figure 3F), as well as significantly lower Ca^2+^ spark frequency (Figure 3G). As a result, TPM myocytes presented a significant reduction in spark-mediated Ca^2+^ leak (Figure 3H). We observed no difference in the SR load between permeabilized myocytes of each genotype (Figure 3I); hence, there was also a significant reduction in the leak normalized to SR load (Figure 3J). We also noted significant differences in the properties of the Ca^2+^ sparks, including full width at half maximum and full duration at half maximum (Figure S5).

Total RyR2-mediated SR Ca^2+^ leak was further quantified in intact isolated cardiomyocytes using the protocol developed by Shannon et al. (Figure 3K),^25^ in which tetracaine is used to block RyR2 in a 0Na^+^/0Ca^2+^ solution. The drop in fluorescence observed during tetracaine application is indicative of the RyR2-mediated Ca^2+^ leak. In basal conditions, SR Ca^2+^ leak was similar in both groups. However, TPM myocytes showed significantly lower Ca^2+^ leak under Iso stimulation, compared to WT myocytes (Figure 3L). This difference was also evident when the leak was normalized to the SR load, measured at the end of the protocol by applying a pulse of caffeine (10 mM) (Figure 3M). CaT amplitude, time to peak and SR load were not different between WT and TPM (Figure S6), consistent with our previous data. Collectively, these results suggest that TPM channels are less active and, therefore, less leaky than WT channels in basal conditions and under Iso stimulation. An important corollary to this data is that that the three canonical phospho-sites contribute to regulate RyR2-mediated Ca^2+^ leak.

### TPM mice are susceptible to cardiac arrhythmia, but cardiomyocytes show fewer spontaneous Ca^2+^ release events

A phospho-mimetic substitution at S2814 (RyR2-S2814D) has been reported to create leakier RyR2 channels that make the heart susceptible to catecholaminergic arrhythmia,^26^ while phospho-ablation (RyR2-S2814A) prevents Ca^2+^ leak and protects against arrhythmia.^13^ We therefore tested the susceptibility of TPM mice to develop catecholaminergic arrhythmia by using a common pharmacological challenge (2 mg/kg epinephrine and 120 mg/kg caffeine, i.p.; Figure 4A). Remarkably, 72.73% (16/22) of TPM mice developed bidirectional ventricular tachycardia, a typical arrhythmia associated with Ca^2+^ mishandling, compared to only 4.5% (1/22) in WT mice (Figure 4B).

**Figure 4.**
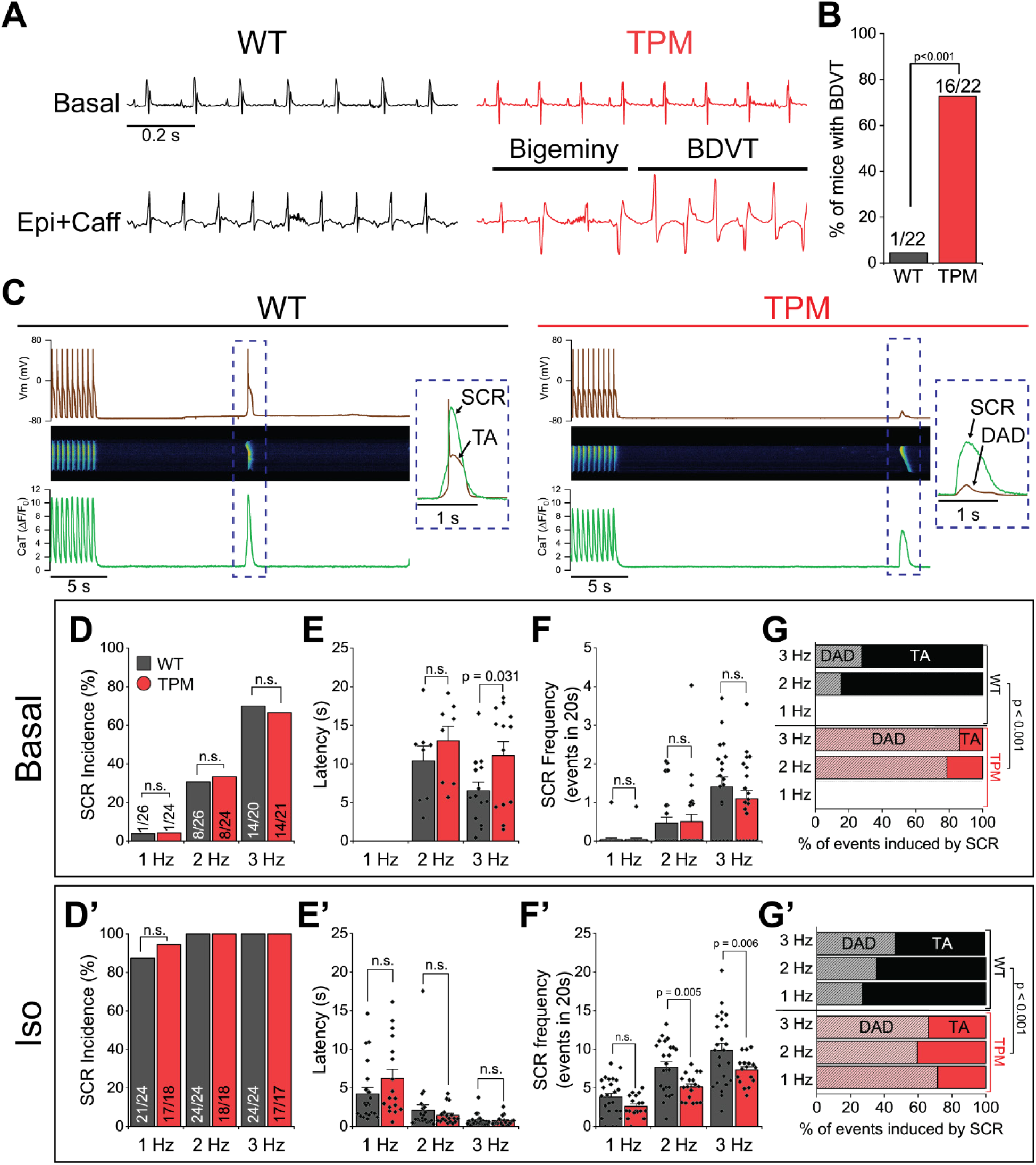
TPM mice have increased susceptibility to arrhythmia despite reduced incidence of diastolic Ca^2+^ release. **A.** Representative ECG traces from anesthetized mice undergoing arrhythmia challenge (Epi+Caff, epinephrine 2 mg/kg, caffeine 120 mg/kg, i.p.). **B.** Incidence of bidirectional ventricular tachycardia (BDVT) during the arrhythmia challenge. TPM mice are significantly more susceptible to BDVT than WT control (n = 22 mice per genotype. Chi-square test). **C.** Representative traces of simultaneous recordings of membrane potential (Vm) and confocal CaTs (Vm/CaT recordings) used to evaluate the incidence of spontaneous Ca release (SCR events), delayed afterdepolarizations (DADs) and triggered activity (TA). Upper panel: current clamp measurements of Vm; middle panel: confocal image of CaTs; lower panel: quantification of fluorescence intensity profile of CaT. Inset shows the superimposed voltage and CaT signal indicated in the dashed blue box. **D,D’.** Incidence of SCR events during a 20 s monitoring period after a train of stimulation at the indicated frequencies. **E,E’.** Latency time to first SCR event following immediately after the last stimulation. TPM myocytes show longer latencies at 3 Hz in basal conditions. **F,F’.** Number of SCR events recorded during 20 s of monitoring. TPM myocytes show fewer SCR events under Iso. **G,G’.** Classification of electrical events induced by SCR into DADs (bumps in the membrane potential) and TA (spontaneous action potentials). Recordings were made under basal conditions (**D-G**) and in the presence of Iso 300 nM (**D’-G’**). Overall, TPM myocytes are less susceptible to SCR than WT controls, while SCR events in TPM myocytes are also less likely to induce TA than in the WT. (N = 13 WT, 7 TPM mice; n = 19-26 WT, 21-24 TPM myocytes [Basal]. N = 10 WT, 11 TPM mice; n = 27 WT, 18 TPM myocytes [Iso]. Chi Square [D-D’,G-G’]; Two-way ANOVA (Hold-Sidak method) [E-E’,F-F’])

To investigate the cellular mechanisms underlying the catecholaminergic arrhythmias in TPM mice, we employed path clamp and confocal microscopy to simultaneously record membrane potential (Vm) and CaTs in isolated cardiomyocytes (Vm/CaT, Figure 4C). This allowed us to evaluate the susceptibility of myocytes to develop spontaneous Ca^2+^ release (SCR) events and their effect on Vm stability. Cells were paced at various rates (1-3 Hz) followed by a 20-second resting period (Figure S7), in basal conditions and under 300 nM Iso stimulation. Interestingly, the incidence of SCR events was not different between genotypes either in basal conditions (Figure 4D) or under Iso stimulation (Figure 4D’). TPM myocytes showed significantly longer latency (time from the last paced CaT to the first SCR event) following 3 Hz pacing under basal conditions (Figure 4E). While the frequency of SCR events was not different in basal conditions (Figure 4F), TPM myocytes showed fewer SCR events under Iso stimulation (Figure 4F’). We categorized SCR-induced disturbances of Vm into delayed after depolarizations (DADs, sub-threshold diastolic depolarizations of Vm) and triggered activity (TA, spontaneous diastolic APs) (Figure 4C). Under both basal conditions and Iso stimulation, SCR events commonly induced DADs in TPM cardiomyocytes, as opposed to TA in WT cells (Figure 4G,G’). Therefore, TPM myocytes exhibit fewer SCRs, and these events generated mainly subthreshold Vm disturbances (DADs). These results are consistent with those presented earlier, showing that RyR2-TPM channels are less leaky than WT channels. They further suggest that diastolic Ca^2+^ release is not the cellular mechanism of the arrhythmias observed in the whole animal.

### TPM myocytes are prone to early afterdepolarizations (EADs)

We further used Vm/CaT recordings to analyze AP properties and assess the incidence of early afterdepolarizations (EADs, depolarizations ≥10 mV during the repolarization phase of the AP) as a potential cellular mechanism for cardiac arrhythmia. Remarkably, WT myocytes showed smooth phase-3 repolarization of the AP, while TPM myocytes commonly displayed a depolarization spike consistent with EAD formation (Figure 5A, Figure S8A). TPM myocytes were more susceptible to EADs at 2-3 Hz in basal conditions and at all frequencies under Iso stimulation (Figure 5B). In basal conditions, TPM myocytes showed EADs that developed progressively at faster pacing rates (Figure S8A). Under Iso, EADs consistently appeared at all pacing frequencies (Figure 5A,B).

**Figure 5.**
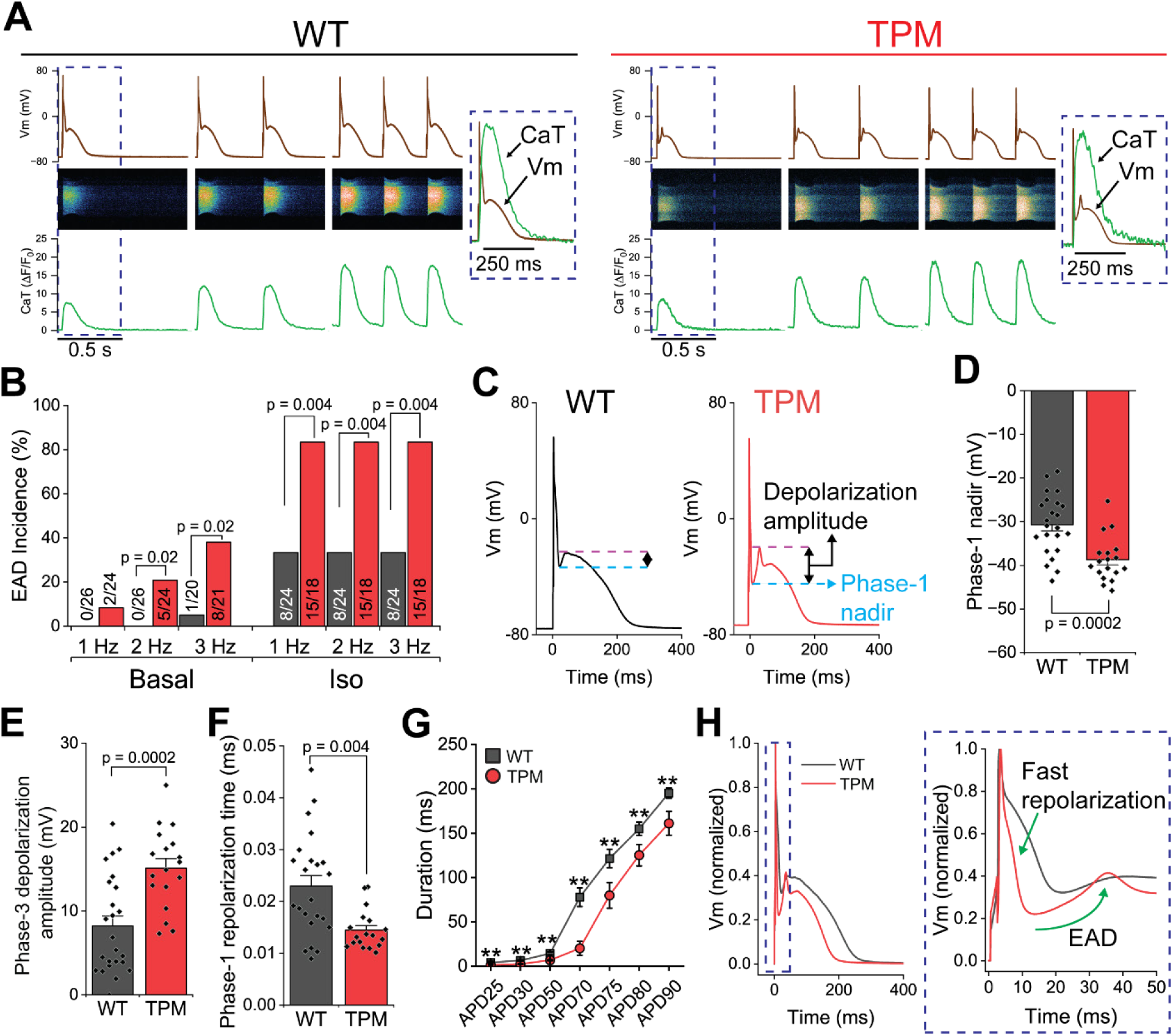
TPM myocytes show faster phase-1, and phase-3 EADs. **A**. Representative Vm/CaT recordings at 1-3 Hz pacing rates under Iso stimulation used to evaluate the incidence of EADs. Left panel: WT; right panel: TPM; brown traces: AP signal; green traces: CaT signal quantified from the confocal images shown. Superimposed Vm/CaT signals are shown in dashed boxes. Under Iso stimulation, EADs were consistently induced at all pacing frequencies in TPM myocytes. **B**. Incidence of cells showing EADs at different pacing rates under basal conditions and Iso stimulation. In basal conditions, the incidence of cells with EADs is significantly higher in TPM cells at 2 Hz and 3 Hz. Upon Iso stimulation, TPM cells had a significantly higher incidence of EADs at all pacing rates. **C-F**. Representative AP morphology under Iso stimulation at 1 Hz pacing (C). TPM cells have a deep notch at the phase-1 nadir followed by a second depolarization upstroke. We quantified the phase-1 nadir potential (D), the amplitude of the second depolarization (E) and the phase-1 repolarization time (F). **G**. Action potential duration under Iso stimulation. TPM cells showed significantly shorter durations from APD20 to APD90 (* p < 0.05, **p < 0.01 TPM vs WT at the same APD). **H**. Superimposed representative AP morphology under Iso stimulation summarizing the findings presented in this figure. TPM myocytes show faster phase-1 that reaches a more negative nadir and exhibit more pronounced depolarization consistent with EAD formation (N = 13 WT, 7 TPM mice; n = 19-26 WT, 21-24 TPM myocytes [Basal]. N = 10 WT, 11 TPM; n = 27 WT, 18 TPM myocytes [Iso]. Chi-square [B]; T test or Mann-Whitney Rank Sum Test [E-G]; two-way ANOVA Holm-Sidak method [H]).

Under basal conditions, WT myocytes exhibited the typical AP shape of murine ventricular myocytes, while some TPM myocytes developed EADs during phase-3 and appeared to have longer AP duration (APD) (Figure S8B-D). However, no significant difference in APD was detected between WT and TPM myocytes, possibly due to the relatively small proportion of cells with EADs (≤30% in TPM, Figure 5B). Interestingly, TPM myocytes with EADs showed a tendency for shorter AP duration times at APD25, APD30, and APD50 compared to WT cells (Figure S8E-F) as a result of the faster phase-1 that precedes EAD occurrence.

Under Iso stimulation, WT myocytes showed a smooth phase-1 and phase-3 of the AP (Figure 5A,C). In contrast, TPM myocytes commonly displayed a fast and more pronounced phase-1, creating a deep notch in the AP profile that was followed by an EAD spike (Figure 5C). Accordingly, TPM myocytes had a significantly more negative Vm potential at phase-1 nadir (Figure 5D), a larger depolarization voltage prior to entering phase-3 (Figure 5E), and a faster phase-1 repolarization time (Figure 5F). Although AP amplitude and resting membrane potential were not different between genotypes (Figure S9), APD was significantly shorter in TPM myocytes across the entire AP under Iso stimulation (Figure 5G). These results suggest that TPM cells undergo faster phase-1 and exhibit phase-3 EADs (Figure 5H).

### Ca^2+^ chelator EGTA eliminates EADs in TPM myocytes

The analysis of AP morphology in TPM cells with EADs suggests these are phase-3 EADs, widely recognized as initiators of ventricular arrhythmia in mouse ventricles. Characteristically, phase-3 EADs require intact SR Ca^2+^ release and typically occur prior to or simultaneously with the peak of the triggered Ca^2+^ transient.^27,28^ This pattern aligns with our observations in TPM cells (Figure 5A). To investigate the role of SR Ca^2+^ release in the formation of EADs in TPM myocytes, we added 10 mM EGTA, a potent Ca^2+^ chelator, to the patch pipette during Vm/CaT recordings. As expected, EGTA turned the CaT into a spike of low amplitude and duration. Most notably, EADs were completely abolished in all cardiomyocytes from both genotypes in the presence of EGTA (Figure 6A,B). EGTA also normalized APD in TPM myocytes (Figure 6C,D), without affecting the resting membrane potential or AP amplitude (Figure S10). These findings suggest that SR Ca^2+^ release may contribute to the faster phase-1, and phase-3 EAD formation in TPM myocytes, even in the absence of overt systolic Ca^2+^ mishandling.

**Figure 6.**
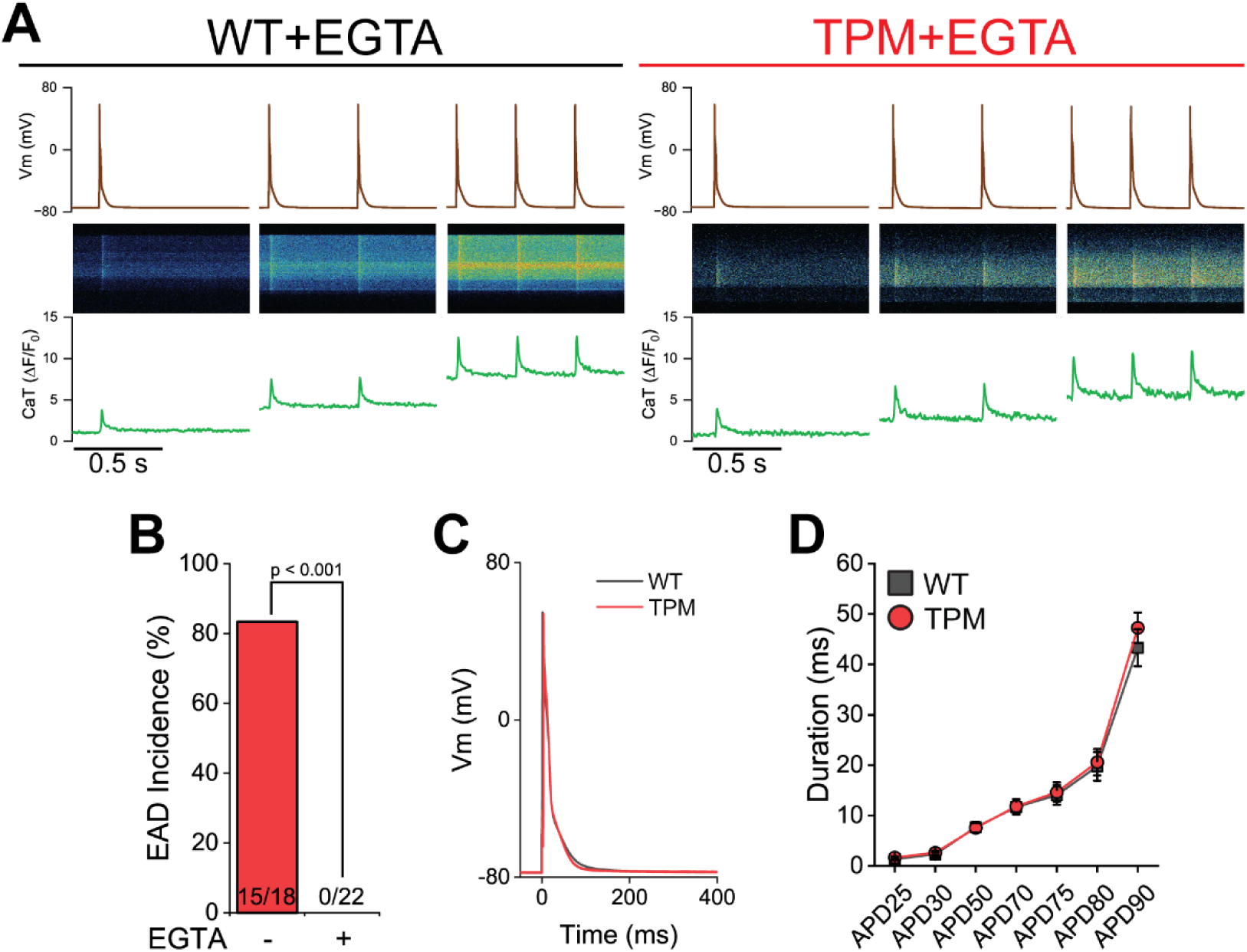
Addition of EGTA to the patch pipette suppresses EAD in TPM myocytes. **A**. Representative traces of Vm/CaT recording with 10 mM EGTA in the patch pipette under 300 nM Iso stimulation at 1-3 Hz pacing. Upper panel: Vm recordings. Middle pannel: confocal line scan image. Bottom panel: CaT profile quantified from the middle panel. EGTA chelates intracellular Ca^2+^, narrowing the AP in both WT and TPM myocytes. **B**. Quantification of EAD incidence in TPM myocytes paced at 1 Hz without EGTA (data from Figure 5B) and with EGTA in the patch pipette. EGTA completely abolished EADs **C-D**. Superimposed APs (C) and quantification of APD (D) in WT and TPM myocytes patched with EGTA. TPM and WT myocytes have identical AP morphology and duration in the presence of EGTA (N = 6 mice per genotype; n = 18 WT, 22 TPM myocytes. Chi-square [B], Mann-Whitney Rank Sum Test or T-test [D]).

### TPM myocytes show reduced *I_CaL_* and increased reactivation of *I_Na_*

Our findings with EGTA indicate the mechanisms generating the fast phase-1, and phase-3 EADs in TPM myocytes may be influenced by Ca^2+^-dependent regulation of sarcolemmal currents. Recognizing the involvement of *I_CaL_* during phase-1 of the mouse cardiac AP,^29^ we hypothesized that the faster phase-1 in TPM cells may be due to a decrease in *I_CaL_*. We therefore measured *I_CaL_* across a range of testing potentials under Iso stimulation (Figure 7A) and compared current density and inactivation kinetics between WT and TPM myocytes (Figure 7B). TPM cells exhibited lower *I_CaL_* density at -10 mV, 10 mV, and 20 mV (Figure 7C), while the inactivation constant of *I_CaL_* (tau fast at -10 mV) was similar between genotypes (Figure 7D). Given that expression of Cav1.2 (the pore-forming subunit of the LTCC) is unchanged in TPM samples (Figure S3), these data indicate that dysregulation of the peak *I_CaL_* that may contribute to the fast and hyperpolarized phase-1 nadir.

**Figure 7.**
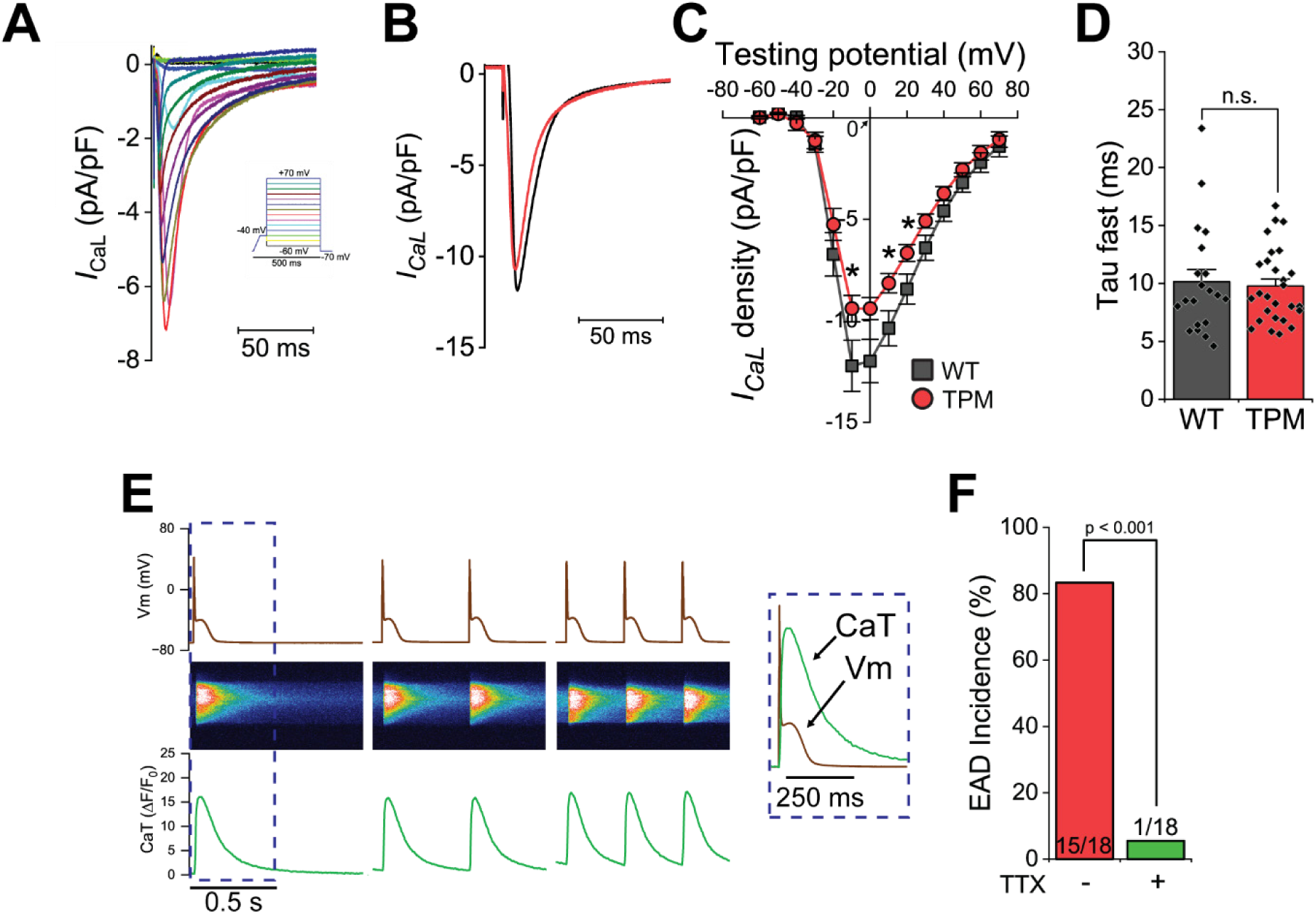
TPM myocytes show reduced *I_CaL_* and non-equilibrium reactivation of *I_Na_*. **A**. Representative *I_CaL_* traces and testing potentials (insert) in the voltage-clamp mode recorded under Iso stimulation. **B**. Representative *I_CaL_* traces at -10 mV from WT and TPM myocytes. **C**. I-V curve of *I_CaL_* density. TPM and WT cells have comparable voltage-dependent activation and inactivation kinetics with maximum activation at -10 mV. However, TPM cells exhibited significantly lower *I_CaL_* density at -10 mV, 10 mV, and 20 mV testing potentials. **D**. Fast decay time of *I_CaL_* inactivation measured by exponential fitting at -10 mV. **E**. Representative AP/CaT recording from a TPM myocyte incubated with TTX under Iso stimulation. Upper: AP traces; Middle: confocal image; lower: fluorescence quantification. Right shows enlarged superimposed AP/CaT signal from the dashed box. **F**. Incidence of EAD in TPM cells under Iso stimulation without TTX (data from Figure 5B) or in the presence of TTX (N = 11 Iso, 5 Iso+TTX TPM mice; n = 18 Iso, 18 Iso+TTX myocytes. Chi-square).

Finally, evaluated the potential ionic current underlying EAD formation. The faster phase-1 reached a negative membrane potential (below -40 mV) that is incompatible with *I_CaL_* reactivation but may accelerate *I_Na_* recovery from inactivation.^28^ We thus hypothesized that the EADs in TPM cells are caused by reactivation of *I_Na_*. To test this idea, we used low concentration tetrodotoxin (TTX, 5 µM) to partially block *I_Na_* under Iso stimulation. At this concentration, TTX mildly reduces peak *I_Na_* but mitigates EADs formation.^28^ TTX treatment significantly reduced the occurrence of EADs in TPM cells, dropping the incidence at 1 Hz pacing from 83.3% to 5.6% (1 out of 18, p<0.001. Chi-square) (Figure 7I). This supports the notion that EADs in TPM cells are caused by non-equilibrium reactivation of *I_Na_*. TTX application also resulted in a lower AP amplitude, more positive phase-1 nadir, lower depolarization prior to phase-3, and a shift from TA to DADs during SCR-induced electrical events (Figure S11).

## DISCUSSION

A classical, widely accepted mechanism for adrenergic regulation of cardiac contractility and Ca^2+^ homeostasis involves increased “trigger Ca^2+^” through LTCC (via PKA-mediated phosphorylation on its inhibitor Rad)^3^ and SR Ca^2+^ uptake by SERCA2 (via PKA-mediated phosphorylation on its inhibitor PLB).^1^ RyR2 is among the first substrates to undergo phosphorylation upon β1-adrenergic stimulation. Yet, the contribution of RyR2 phosphorylation to achieving a full β1-adrenergic response remains controversial, ranging from a central mechanism^8^ to functionally inconsequential.^11^ Our focus in this study was on the three canonical sites (S2031, S2808 and S2814) because they are the sites known to be phosphorylated during adrenergic stimulation: PKA phosphorylates S2031 and S2808, while CaMKII phosphorylates S2808 and S2814.^6^ S2368 (phosphorylated by SPEG)^30^ and T2810 (a hypothesized PKC site)^31^ are other identified phosphorylation sites that are not linked to adrenergic regulation of the channel and, therefore, fall beyond the scope of this work. We generated the RyR2-S2031A/S2808A/S2814A (triple phosphomutant, TPM) mouse model to evaluate the role of the three canonical sites in the regulation of the channel. We speculated that any difference in the cardiac adrenergic response between TPM and WT mice and myocytes would stem from the regulatory effect that phosphorylation of these sites may exert on RyR2.

### Is RyR2 phosphorylation essential for the β-adrenergic response?

Our data show that TPM mice have nearly identical chronotropic and inotropic responses to adrenergic stimulation compared to WT controls at the whole-animal and whole-myocyte levels (Figure 2). These are remarkable findings that suggest the three canonical phosphorylation sites in RyR2 are not be crucial for the global adrenergic response. These results should not be misconstrued as supportive of the idea that RyR2 phosphorylation doesn’t play a role in the β-adrenergic response. In this view, RyR2 would act as a passive gatekeeper, activated *I_CaL_* and allowing Ca^2+^ follow an electrochemical gradient, leaving no role for direct regulation of RyR2 open probability or Ca^2+^ sensitivity.^32^ However, such simplified scheme has a conundrum: if RyR2 is only a passive gate, why does the channel keep an intricate regulatory machinery, with built-in kinases and phosphatases, and undergoing evident phosphorylation upon adrenergic stimulation?

To address this question, it is critical to first establish a distinction between the adrenergic responses of the whole heart and whole cardiomyocyte, and that detected at the subcellular level. The former involves positive chronotropy (regulated at the sinus node), as well as positive inotropy and lusitropy (regulated in the ventricles).^33^ Despite previous reports to the contrary, as we have discussed elsewhere,^7^ data consistently show that preventing phosphorylation of RyR2 at individual canonical sites does not hinder inotropy or chronotropy. For instance, we showed that S2808A mice^10,11^ and S2808/S2814A double knock-in mice exhibit normal adrenergic response.^17^ Nonetheless, in these studies we could not reject that the availability of one or more remaining canonical sites could account for the global adrenergic response. Here, we demonstrate that removal of all three sites leads to the same outcome.

The literature and our data consistently show that phosphorylation of RyR2 at the canonical sites regulates certain subcellular properties of Ca^2+^ homeostasis during adrenergic stimulation.^34^ We observed this in the TPM mouse and other phospho-deficient models, in which Iso stimulation does not fully elicit its expected effects on Ca^2+^ sparks properties, SR Ca^2+^ leak and Ca^2+^ waves.^15,35^ Our findings show that the inability of TPM channels to undergo phosphorylation at the canonical sites prevents RyR2 from increasing SR Ca^2+^ leak upon adrenergic stimulation. The reduced Ca^2+^ leak, only ∼3% of the total SR load (Figure 3D), may have minimum impact on global Ca^2+^ dynamics and systolic Ca^2+^ release, but could significantly affect the balance of sarcolemmal currents. This may be particularly relevant at the dyad, where t-tubules form tight couplings with the junctional SR. Furthermore, phosphorylation of RyR2 has been shown to participate in the rearrangement of channels clusters.^36–38^ It is therefore plausible that, even if RyR2 phosphorylation at the canonical sites is not required for the global adrenergic response (chronotropy and inotropy), it is essential to regulate RyR2 organization and finetune subcellular Ca^2+^ release, which may have important pathophysiological implications.

These results don’t negate the potential significance of other, yet-to-be-identified phosphorylation sites. Indeed, we detected significant incorporation of ^32^P into TPM channels by PKA and CaMKII (Figure 1). S2808 was initially proposed as the only PKA site in RyR2, in part because this enzyme was purportedly unable to phosphorylate S2808A channels,^9^ but these findings are not universally accepted.^20,21^ S2031 was later identified as a second major PKA site with wide dynamic range, given its low basal phosphorylation level (Table S1).^6,20^ It was also reported that the recombinant RyR2 with the double mutation S2808A/S02031A could still undergo PKA-mediated phosphorylation.^20^ Our data using TPM channels are fully consistent with these results: we observed a ∼70% reduction in the amount of ^32^P incorporated by PKA into TPM channels. We surmise that S2031 and S2808 are the major but not the only PKA sites. Similarly, the prevalent idea that S2814 is the unique CaMKII phosphorylation site arose from a report showing that S2814A channels cannot be phosphorylated by this enzyme.^12^ This is inconsistent with previous data and our present findings. Early biochemical studies demonstrated that CaMKII is able to incorporate ∼4x more ^32^P into RyR2 than PKA.^23^ If there are two already known unambiguous PKA sites (S2808 and S2031) and one of them has very low basal phosphorylation (S2031), then it is unlikely that a single CaMKII site (S2814) accounts for such large difference in ^32^P incorporation. S2808 is also phosphorylated by CaMKII,^6^ further undermining the notion of a single CaMKII residue. Consistent with these results, we determined that CaMKII introduced nearly the same amount of ^32^P into WT and TPM channels. While the canonical sites may regulate subcellular Ca^2+^ handling during the adrenergic response (which is impaired in TPM mice), our results still leave the open possibility that other unknown sites could participate in chronotropy and inotropy (which are normal in the TPM).

### Arrhythmia mechanism in TPM mice: the double-edged sword of SR Ca^2+^ leak

TPM mice are highly susceptible to cardiac arrhythmia under catecholaminergic stimulation (Figure 4A-B), a finding that is not entirely surprising given that single point mutations that alter the biophysical properties of RyR2 can lead to this phenotype. Nearly two hundred mutations in the gene encoding the channel have been linked to two inherited arrhythmia disorders in human patients: catecholaminergic polymorphic ventricular tachycardia (CPVT) due to gain-of-function mutations^39^ and Ca^2+^ release deficiency syndrome (CRDS) due to loss-of-function mutations.^40^ TPM channels, however, display unique biophysical alterations that suggest a complex mechanism of arrhythmia arising from the inability of TPM to undergo phosphorylation at the canonical sites.

The type of arrhythmia present in TPM mice, bidirectional ventricular tachycardia under adrenergic stress, which is typical of CPVT. Yet, TPM cardiomyocytes do not display the cellular hallmarks of CPVT. Instead, we found fewer diastolic SCR events and DADs (Figure 4C-G). Based on the lower SR Ca^2+^ leak under similar SR load (Figure 3E-M), our data show that TPM channels display a constitutive loss-of-function. Interestingly, a cellular mechanism by which a loss-of-function mutation leads to cardiac arrhythmia was through EADs,^41^ similar to our observations in the TPM model (Figure 5). RyR2 loss-of-function is linked to CRDS, but mice harboring such mutations are resistant to arrhythmias under adrenergic stress.^40^ It may seem paradoxical that TPM mice display a mixed phenotype between CRDS and CPVT, but bidirectional tachycardia induced by EADs (rather than DADs) has been reported in other mouse models of arrhythmia.^42^

TPM channels are characterized by their ability to release Ca^2+^ at normal levels during ECC; however, they exhibit a subtle but constant loss-of-function that impacts diastolic SR Ca^2+^ leak. The delicate balance between Ca^2+^ release and Ca^2+^ leak is critical to maintain normal CaTs, and alterations in any of these processes may destabilize Ca^2+^ homeostasis, especially during adrenergic stimulation. In this scenario, SR Ca^2+^ leak becomes double-edged sword because too much or too little leak are detrimental to cardiac function. Increased SR Ca^2+^ leak a well-known culprit of cardiac arrhythmia,^19^ and here we show in the TPM model that a discrete amount of leak may also be necessary to maintain Ca^2+^ homeostasis. Hence, WT RyR2 channels, which can undergo phosphorylation at the three canonical sites, increase their sensitivity to activation by Ca^2+^ during adrenergic stimulation, leading to a moderate increase in Ca^2+^ leak that prevents SR Ca^2+^ overload. TPM RyR2 channels, on the other hand, lack such critical dynamic regulation, which manifests as increased propensity to arrhythmia through EADs. Therefore, phosphorylation of the three canonical sites may be critical to the physiological tuning of SR Ca^2+^ leak.

While we did not observe overt Ca^2+^ mishandling during pacing, the addition of EGTA to the patch pipette completely abolished the APD alterations and EADs in TPM cardiomyocytes (Figure 6). This key piece of data strongly suggests that SR Ca^2+^ release is required for EAD formation. We observed faster AP phase-1 prior to the EADs (Figure 5) and a mild but significant reduction in peak *I_CaL_* (Figure 7). In mouse cardiomyocytes, *I_CaL_* largely occurs during phase-1 of the AP. We hypothesize that a reduction in *I_CaL_* enhances repolarization and allows the Vm to reach a more negative phase-1 nadir (Figure 5).^29^ This negative Vm favors recovery of Nav1.5 from inactivation, priming it for reactivation. Indeed, the application of low TTX concentrations during AP recordings significantly reduced the incidence of EADs (Figure 7H-I). This suggests that the EADs are the result of non-equilibrium reactivation of *I*_Na_, as has been suggested previously in mouse ventricular myocytes.^28^

### Conclusions

An early model of RyR2 regulation by phosphorylation where one kinase phosphorylates one site to produce one well-defined effect^43^ appears now oversimplified. This model has proven insufficient to reconcile results from recent studies,^7^ including those highlighting the third canonical site, S2031. Our data also strongly suggest that S2808 and S2814 may no longer be considered exclusive PKA and CaMKII phosphorylation sites, respectively. Phosphorylation of RyR2 increases channel activity, but it has also been proposed that dephosphorylation using protein phosphatases leads to a similar outcome.^44–46^ These studies took an agnostic approach that did not look at specific phospho-sites, but rather evaluated the overall phosphorylation state of the channel. We continue to propose that a model of RyR2 regulation by phosphorylation/dephosphorylation must consider the complex interplay between phosphorylation sites and their potential hierarchy in the regulation of this large ion channel.^43^ Our study adds critical information to continue building this model.

Altogether, this study provides strong experimental evidence to propose that the canonical phosphorylation sites in RyR2 (1) are not critical for the global adrenergic response in the heart (mainly inotropy and chronotropy), but (2) they are required to dynamically regulate a discrete physiological Ca^2+^ leak during β-adrenergic stimulation that contributes to the electrical stability of cardiomyocytes. Future studies will need to elucidate the determinants of the interplay between SR Ca^2+^ leak and the regulation of sarcolemmal Ca^2+^ currents.

## ACKNOWLEDGEMENTS

We thank the University of Wisconsin-Madison Biotechnology Center Animal Models Core and Advanced Genome Editing Laboratory (RRID:SCR_021070) for their contributions in generating genome edited models.

## Non-standard Abbreviations and Acronyms

RyR2: Ryanodine receptor 2
ECC: excitation-contraction coupling
AP: action potential
LTCC: L-type Ca^2+^ channels
*I_CaL_*: inward L-type Ca^2+^ current
SR: sarcoplasmic reticulum
CICR: Ca^2+^-induced Ca^2+^ release
CaT: Ca2+ transient
Vm: transmembrane potential
SERCA2: SR Ca^2+^ pump
NCX: Na^+^/Ca^2+^ exchanger
β1-Ars: β1-adrenergic receptors
PKA: protein kinase A
CaMKII: Ca^2+^/calmodulin-dependent kinase II
PLB: phospholamban
TPM: triple phospho-mutant
DAD: delayed after-depolarization
AP: action potential
EAD: early after-depolarizations
Vm/CaT: simultaneous membrane potential and Ca^2+^ transient recordings
WBs: western blots
Iso: isoproterenol
Vm: membrane potential
SCR: spontaneous Ca^2+^ release
TA: triggered activity
APD: action potential duration
*I_Na_*: Na^+^ current
TTX: tetrodotoxin

